# Development of an HK-fbp1-based vaccine platform against *Cryptococcus neoformans*

**DOI:** 10.64898/2026.10.21.760669

**Authors:** Yu Zhang, Yina Wang, Keyi Wang, Lu Gan, Aaron Lemmer, Siddhi Pawar, Amariliz Rivera, Chaoyang Xue

**Affiliations:** Public Health Research Institute, and Department of Microbiology, Biochemistry and Molecular Genetics, New Jersey Medical School, Rutgers University, Newark, NJ, 07103, United States; Center for Immunity and Inflammation, and Department of Pediatrics, New Jersey Medical School, Rutgers University, Newark, NJ, 07103, United States

**Author notes:** **Corresponding author:** Chaoyang Xue, The Public Health Research Institute Rutgers University ICPH building Room W210E, 225 Warren Street, Newark, NJ 07103.

**Keywords:** *Cryptococcus neoformans*, heat-killed fungal vaccine, *fbp1*Δ mutant, CpG ODN 2395, intramuscular immunization, protective immunity, antifungal vaccine, T cell immunity

## Abstract

Cryptococcal meningitis remains a leading cause of death among HIV/AIDS patients, and no antifungal vaccine is available in clinical use. We previously showed that mice immunized with a heat-killed preparation of *Cryptococcus neoformans* (*Cn*) *fbp1*Δ mutant (HK-fbp1) are protected against cryptococcosis and other invasive mycoses, suggesting that HK-fbp1 is an intriguing vaccine candidate for further study. Recent studies have reported that heat-killed wild type H99 (HK-H99) can also offer some protection in other vaccination models. Here we set out to decipher the differential protective capacity of HK-fbp1 versus HK-H99 across multiple conditions. We observed that HK-fbp1 confers superior protection in both prophylactic and therapeutic intranasal immunization settings than HK-H99, which was associated with greater activation of pulmonary CD4⁺ T cells and stronger airway IFN-*γ* production. In addition, HK-fbp1 preparations were highly stable, retaining full protective efficacy even after 5 years of frozen storage. Mice intranasally immunized with two-dose HK-fbp1 were protected against challenge by up to 10⁶ H99 cells, a 100-fold increase over the commonly used lethal inoculum. Addition of a CpG adjuvant ODN to HK-fbp1 (HK-bp1/ODN) led to a 5-fold reduction of the intranasal vaccine inoculum required for protection, and itssingle-dose intramuscular vaccination induced protection persisted for at least 3 months post vaccination. Together, these findings highlight the robust protective capacity and formulation versatility of HK-fbp1, supporting its further development as a stable, dose-sparing antifungal vaccine candidate.

**Importance:** Vaccines against viral and bacterial infections have had a transformative impact in our fight against infectious diseases. However, there are no fungal vaccines currently approved for clinical use. This study provides a full characterization of a potent whole cell fungal vaccine we recently developed based on heat-killed *Cryptococcus neoformans fbp1*Δ mutant (HK-fbp1) and its adjuvanted version. Our data demonstrates that HK-fbp1 and its adjuvanted version have superior protection advantage than heat-killed wild type H99. Importantly, the HK-fbp1 is highly stable with its preparation remained protective potency after years of frozen storage. Including a CpG adjuvant ODN further reduced the antigen requirement in the intranasal route, and provided HK-fbp1 a highly durable protection after single intramuscular vaccination. These clinically relevant attributes of a successful vaccine, including high stability, dose-sparing potential, and durable efficacy following a single dose, strengthen the rationale for developing HK-fbp1 as a versatile whole-cell antifungal platform.

## Introduction

Invasive fungal infections are a major cause of death in immunocompromised populations (1, 2). Among them, *Cryptococcus neoformans* is the leading cause of life-threatening fungal meningoencephalitis, particularly in individuals with impaired cell-mediated immunity. Despite advances in antifungal treatment, cryptococcal meningitis remains a major cause of mortality among people with advanced HIV disease, accounting for an estimated 19% of AIDS-related deaths worldwide (3). The absence of a licensed vaccine against human invasive fungal infections further underscores the need for effective preventive strategies (4-6). Whole-cell vaccines represent a promising approach to fungal vaccine development because they preserve a broad repertoire of pathogen-derived antigens and immunostimulatory components capable of engaging both innate and adaptive immunity. In particular, heat-inactivated formulations eliminate the risk of vaccine-strain replication, an important consideration for immunocompromised populations. Several genetically engineered *C. neoformans* strains have demonstrated protective potential in experimental vaccination models, including the IFN-γ-expressing H99γ strain, the sterylglucosidase-deficient *sgl1*Δ mutant, the *ZNF2*-overexpressing strain, the chitin deacetylase-deficient *cda1*Δ *cda2*Δ *cda3*Δ mutant, and the F-box protein-deficient *fbp1*Δ mutant (7-15). However, protective efficacy is not necessarily preserved following heat inactivation. For example, the strong protection induced by live H99γ is lost after heat killing (7), whereas other cryptococcal mutants retain their protective capacity in inactivated formulations (9-15). These observations suggest that vaccine efficacy depends not simply on the presence of fungal antigen amount but also on the immunogenic properties of the vaccine strain.

Among these candidates, heat-killed *fbp1*Δ cells (HK-fbp1) have demonstrated strong protective efficacy following intranasal immunization. Previous studies showed that HK-fbp1 protects mice against lethal pulmonary challenge with *C. neoformans* and *C. gattii*, provides cross-protection against *Aspergillus fumigatus*, and confers partial protection against *Candida albicans* (12, 13). This protection requires IFN-γ signaling but can be maintained in the absence of CD4⁺ T cells under certain vaccination conditions through compensatory CD8⁺ T-cell responses (13, 14). Moreover, early therapeutic administration of HK-fbp1 improves survival and restricts fungal dissemination in mice with established cryptococcal infection (14). Together, these findings establish HK-fbp1 as a promising antifungal vaccine candidate with both prophylactic and therapeutic potential. Despite these encouraging findings, several aspects of HK-fbp1 vaccination remain insufficiently characterized. Although protection is dose dependent (13), the ability of HK-fbp1 to withstand increasingly severe infectious challenges and retain its protective efficacy after prolonged storage has not been fully established. Furthermore, the extent to which its superior immunogenicity translates into an advantage over heat-killed wild-type cells remains unclear. Previous studies have shown that heat-killed wild-type H99 (HK-H99) can provide measurable protection when administered at sufficiently high doses (16, 17), and such whole cell induced protective immunity is associated with its culture conditions (18), raising the possibility that the protective advantage of HK-fbp1 depends on vaccine dose. Hence, a systematic comparison analysis between HK-fbp1 and HK-H99 is needed to gain a better understanding of the HK-fbp1 vaccine mediated protection. Finally, the established intranasal vaccination strategy requires relatively high antigen doses and repeated administration, presenting practical challenges for further vaccine development. Adjuvantation with CpG oligodeoxynucleotides (ODN) 2395, which activate Toll-like receptor 9 (TLR9) and promote Th1-associated immune responses, offers a potential strategy to improve vaccine efficiency and enable alternative administration routes (19) (Y. Zhang et al., submitted for publication).

In this study, we directly compared the efficacy of HK-fbp1 vaccine with that of HK-H99 under different conditions, systematically evaluated the protective range and long-term stability of HK-fbp1 vaccine. The comparison revealed a protective advantage of HK-fbp1 accompanied by a stronger Th1 response. HK-fbp1 retains protective efficacy after prolonged frozen storage and protects against infectious challenges substantially exceeding the commonly used lethal inoculum. Addition of the CpG adjuvant ODN to HK-fbp1reduces the intranasal vaccine inoculum required for protection and promoted durable protection after a single intramuscular immunization. These findings further define the protective properties of HK-fbp1 and provide a foundation for developing a more practical inactivated antifungal vaccine.

## Results

### HK-fbp1 vaccine confers superior protection and more effective fungal control than heat-killed wild type H99

Whole-cell inactivated vaccines preserve a broad repertoire of pathogen-derived antigens and immunostimulatory components, offering the potential to induce immune responses against multiple targets. In previous studies, we developed an intranasal vaccine strategy based on an *fbp1*Δ mutant of *Cryptococcus neoformans*. Intranasal immunization with the heat-killed preparation (HK-fbp1) conferred robust protection against both *C. neoformans* and *C. gattii* (13). Notably, HK-fbp1 also fully protected mice from a lethal *Aspergillus fumigatus* challenge and prolonged survival following *Candida albicans* and *Staphylococcus aureus* infection, indicating that this vaccine immunization confers broad-spectrum anti-microbial protection (13). In our standard protocol, HK-fbp1 was freshly prepared on the day of immunization to preserve viability-independent antigenicity and to avoid potential loss of potency during storage (Fig. 1A). To achieve maximal protection, mice received two intranasal immunizations with 5 × 10⁷ HK-fbp1 cells 4 weeks apart, and challenged intranasally with 1 × 10^4^ virulent wild-type H99 cells 2 weeks after the boost (Fig. 1B). Whether the robust protection conferred by HK-fbp1 reflects a general protective role of heat-killed *C. neoformans* cells or depends specifically on the *fbp1*Δ mutation had not been carefully tested. We therefore compared the protective efficacy of heat-killed wild-type H99 (HK-H99) directly with that of HK-fbp1 under a number of immunization conditions.

**FIG 1.**
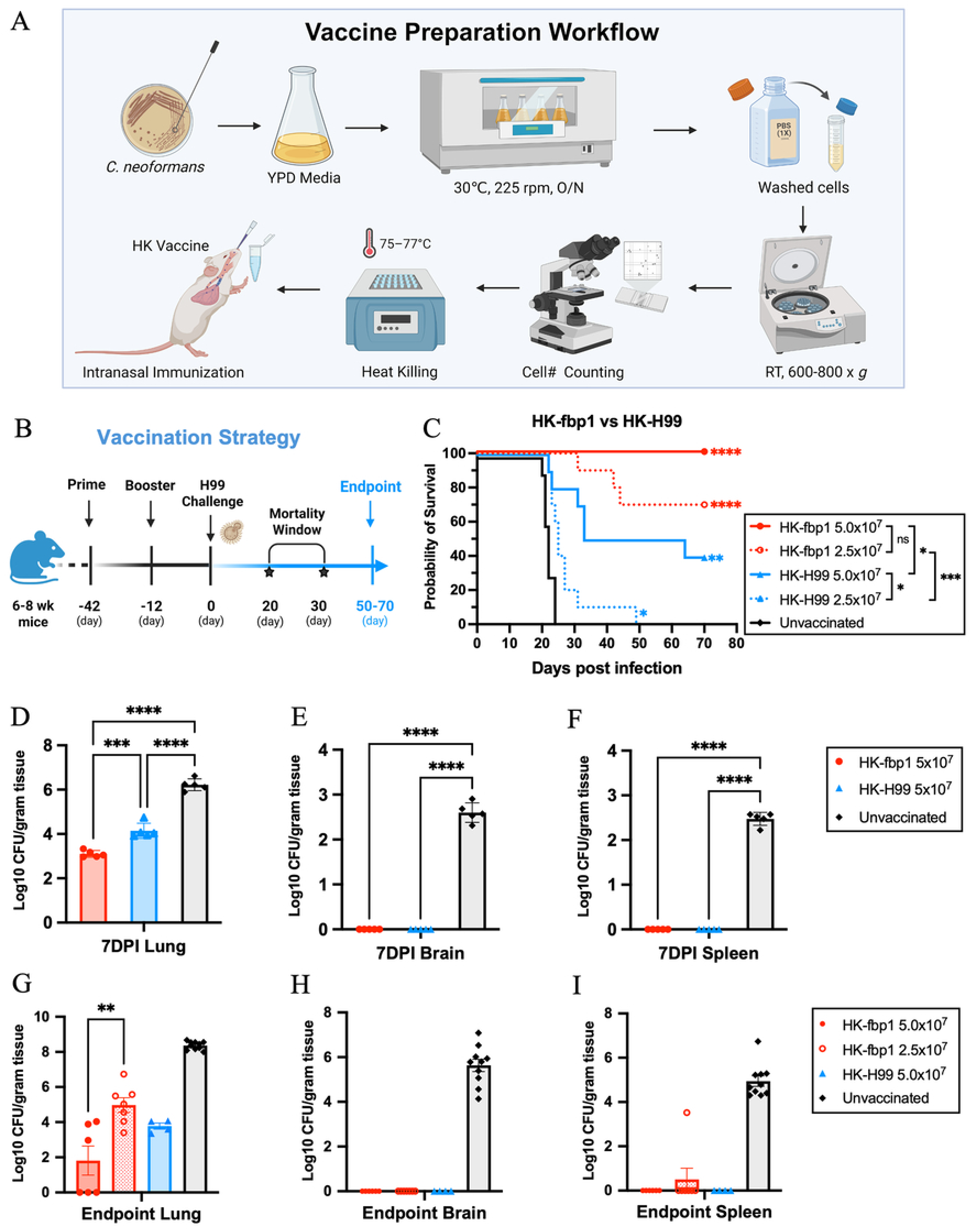
HK-fbp1 confers superior protection and more effective early fungal control than heat-killed wild-type H99. (A) Workflow for preparation of the heat-killed (HK) vaccine. Detailed description is provided in Method. (B) Immunization and challenge schedule. Six-to 8-week-old mice received a priming dose at day −42 and a booster dose at day −12 and were challenged with *C. neoformans* H99 at day 0. Animals were monitored daily for survival, with most deaths in unvaccinated mice occurring between 20 and 30 days post infection (DPI). Surviving animals were monitored to the experimental endpoint at 50 to 70 DPI. The schematic was created with BioRender (A-B). (C) Survival curves of mice vaccinated twice with 5 × 10⁷ or 2.5 × 10⁷ HK-fbp1 or HK-H99 cells and challenged by infection with 10⁴ H99 cells. Vaccination and challenge were performed as described for Fig. 1B. Unvaccinated mice are shown as a control. Six BALB/c mice were used for the HK-fbp1 5 × 10⁷ group, and 10 mice were used for each of the other groups. *, P < 0.05; **, P < 0.01; ****, P < 0.0001 (determined by log rank Mantel-Cox test). Comparisons without an asterisk are not significant. (D to F) Fungal burden in the lungs (D), brains (E), and spleens (F) of a separate cohort of mice at 7 DPI (n = 5 per group). (G to I) Fungal burden in the lungs (G), brains (H), and spleens (I) of the surviving mice at the end of the experiment (n = 4 to 10 per group). The fungal burden of unvaccinated mice at the humane endpoint is shown as a control. **, P < 0.01; ***, P < 0.001; ****, P < 0.0001 (determined by one-way ANOVA with Tukey’s multiple-comparison test within each tissue). Comparisons without an asterisk are not significant.

Mice were immunized intranasally twice with either 5 × 10^7^ or 2.5 × 10^7^ heat-killed cells per dose and challenged as described for Fig. 1B. HK-fbp1 conferred complete protection at the higher vaccine dose and protected 70% of mice at the lower dose (Fig. 1C). HK-H99 was substantially less protective at both doses: 40% of mice survived immunization with 5 × 10^7^ cells, whereas the lower dose vaccination prolonged survival but yielded no endpoint survivors (Fig. 1C). Heat-killed wild-type cells are therefore not inert, but the protection is weaker and requires a higher antigen dose, thus suggesting that HK-fbp1is more immunogenic than HK-H99. Fungal burdens measured at 7 days post infection (7 DPI) showed that immunization with either preparation reduced pulmonary colonization relative to unvaccinated mice and prevented dissemination to the brain and spleen, where no colonies were recovered from any immunized animal (Fig. 1D-F). Pulmonary fungal burdens were significantly higher in mice immunized with 5 x 10^7^ HK-H99 than in HK-fbp1 immunized mice (Fig. 1D), indicating that the two preparations differed in the efficiency of early fungal control even though both prevented extrapulmonary spread. At the study endpoint, surviving mice from both groups retained low-to-moderate pulmonary burdens, with few or no colonies recovered from the brain or spleen (Fig. 1G-I). Among HK-fbp1 vaccinated survivors, residual pulmonary burdens were higher in the lower-dose (2.5 x 10^7^) group than in the higher-dose (5 x 10^7^) group, consistent with dose-dependent control of pulmonary infection (Fig. 1G). Pulmonary fungal burdens were comparable between high-dose HK-fbp1 and high-dose HK-H99 survivors, indicating that within the subset of animals that established protective immunity, both immunizations restricted pulmonary fungal growth to a similar extent. We previously showed that HK-fbp1 is effective not only as a prophylactic vaccine but also as a therapeutic agent in mice already infected with *C. neoformans*, with intervention at 3 DPI rescuing most animals (14). We thus set out to test whether HK-H99 has therapeutic activity as compared to HK-fbp1. Mice were infected intranasally with 10^4^ live H99 cells and treated once at 3 DPI with 5 × 10^7^ HK-fbp1 or HK-H99 cells, respectively, delivered intranasally as previously described (Fig. S1A) (14). HK-fbp1 treatment protected a larger proportion of infected mice to the study endpoint than HK-H99, although the difference between the two treatments did not reach statistical significance (Fig. S1B). Pulmonary fungal burdens in both treated groups were lower than in untreated controls, but dissemination to the brain and spleen was more pronounced in HK-H99 treated mice (Fig. S1C-E).

Together, our findings indicate that heat-killed wild-type cells retain partial activity but require a higher antigen dose and provide less effective early fungal control as compared to HK-fbp1. Under the experimental settings tested, we observed a consistent protective advantage of HK-fbp1 vaccination in both prophylactic and therapeutic settings.

### Th1 polarization and T cell activation distinguish HK-fbp1 from HK-H99 induced lung immunity

Our previous studies have demonstrated that T cells and IFN-*γ* are crucial effectors in vaccine-induced immunity (13, 14) (Y. Zhang et al., submitted for publication). We therefore asked whether the greater protection conferred by HK-fbp1 is associated with a quantitative and qualitatively different T cell response, by profiling mice immunized intranasally with HK-fbp1 or HK-H99 at 7 DPI. We first examined T cell recruitment and activation in the infected lung. At 7 DPI, CD4⁺ T cell recruitment was higher in HK-fbp1 immunized mice than in HK-H99 group (Fig. 2A), and CD8⁺ T cell recruitment showed no difference (Fig. 2B). In CD4⁺ T cell subsets, a greater fraction of recruited cells retained an activated effector phenotype (CD44^hi^CD62L^lo^) in the HK-fbp1 group (Fig. 2C and D), indicating that a larger proportion of CD4⁺ T cells had undergone antigen-driven activation by HK-fbp1 immunization than by HK-H99. In CD8⁺ T cell subsets, a greater percentage showed an activated effector phenotype in the HK-fbp1 group whereas the activated cell number was not significantly different between HK-fbp1 and HK-H99 immunization (Fig. 2E and F). In bronchoalveolar lavage fluid (BALF) collected at 7 DPI, HK-fbp1 immunized mice showed a higher proportion of IFN-γ-producing CD4⁺ T cells and a lower proportion of IL-13-producing CD4⁺ T cells than HK-H99 immunized mice (Fig. 2E and F). Consistent with this, eosinophil numbers in the lung were higher in the HK-H99 group (Fig. S2A), whereas other innate cell populations did not differ detectably between the two groups at this time point (Fig. S2B-E). Together these data indicate that both immunizations favor a type 1-skewed airway response, with HK-fbp1 inducing a stronger IFN-γ response and a smaller IL-13 production than HK-H99. Thus, relative to HK-fbp1, immunization with HK-H99 resulted in less CD4⁺ T cell recruitment, a lower degree of effector T cell phenotype, and reduced Th1 cytokine production, alongside a more pronounced type 2 response, which we consider as a combination less suited to controlling cryptococcal infection.

**FIG 2.**
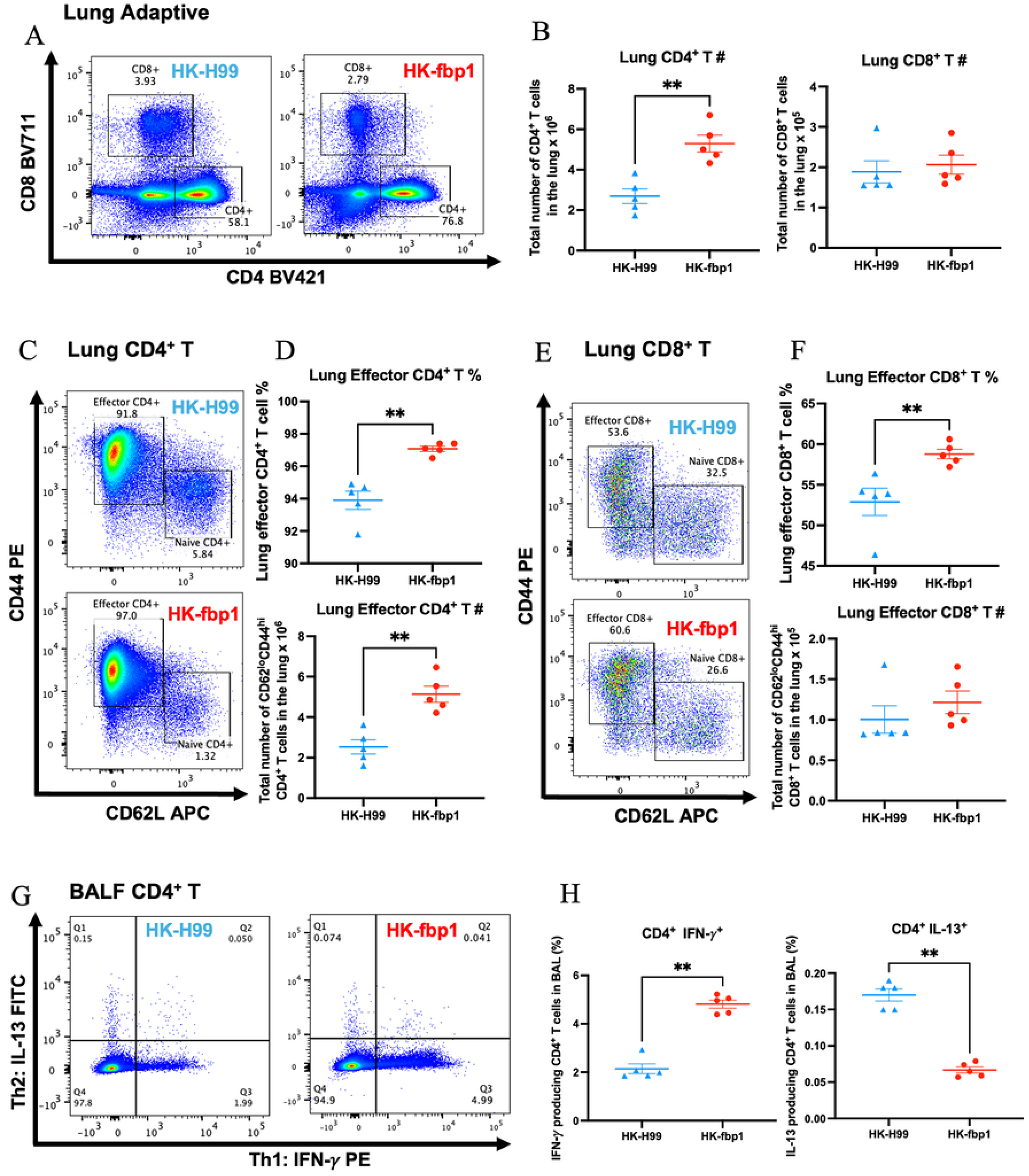
HK-fbp1 elicits a stronger and more activated pulmonary T cell response than HK-H99. Mice were vaccinated twice with 5 × 10⁷ HK-fbp1 or HK-H99 cells and challenged by infection with 10⁴ wild-type H99 cells; BALF and lungs were collected at 7 DPI. Five BALB/c mice were used for each group. (A and B) Representative flow cytometry plots (A) and recruitment (B) of CD4⁺ and CD8⁺ T cells to the lung. (C to F) Representative flow cytometry plots (C, E) and populations (D, F) of lung-recruited CD4⁺ and CD8⁺ T cells with an effector memory (CD44^hi^ CD62L^lo^) phenotype. (G and H) Representative flow cytometry plots (G) and proportions (F) of IFN-γ–producing and IL-13–producing CD4⁺ T cells in BALF. **, P < 0.01 (determined by Mann-Whitney test), comparisons without an asterisk are not significant.

### Intranasal HK-fbp1 confers protection across a broad range of challenge doses and retains efficacy after long-term storage

To define the upper limit of vaccine-induced protection, mice unimmunized or immunized twice with 5 × 10^7^ HK-fbp1 were challenged intranasally with escalating doses of H99 inoculum (10^4^ to 10^7^ CFU). Survival data for all groups are summarized in Table S1. All unvaccinated mice succumbed to every challenge dose, and their median survival shortened progressively as the inoculum increased (Table S1), confirming a dose-dependent acceleration of disease. Intranasal HK-fbp1 immunization protected the majority of mice against challenge with 10^4^ to 10^6^ CFU, with 50% to 86% surviving to the study endpoint, respectively (Fig. 3A and Table S1). Protection was largely overwhelmed at 10^7^ CFU, where only 25% immunized mice survived. Notably, median survival of succumbed mice in this group was identical to that of unvaccinated 10⁷ controls (11 days) (Fig. 3A and Table S1), indicating that at the highest inoculum the vaccine did not slow disease progression in mice that succumbed but instead protected a subset of animals completely. At the humane endpoint, unvaccinated mice harbored comparable fungal burdens in the lung, brain, and spleen irrespective of the challenge dose received (Fig. 3B-D), indicating that death occurred at a consistent terminal burden. In contrast, immunized mice that survived to the study endpoint carried similarly low pulmonary burdens across all challenge doses, and no H99 cells were recovered from the brain or spleen of challenged mice. We next asked whether HK-fbp1 retains protective efficacy after long-term storage. Mice immunized with HK-fbp1 preparations stored at −20°C for 1, 3, or 5 years each remained completely protected against intranasal challenge with 10⁴ H99 cells, comparable to that of freshly prepared vaccine (Fig. 3E). Fungal burdens in the lung, brain, and spleen were comparably reduced across all storage groups and in mice receiving freshly prepared vaccine (Fig. 3F). Together, these results demonstrate that HK-fbp1 remains stable during prolonged frozen storage, supporting its consistent use in future studies. Together, these findings define the protective range of intranasal HK-fbp1 vaccination, demonstrating substantial protection against a broad range of H99 challenge doses, although efficacy declines at the inoculums higher than 10^6^ CFU. Importantly, vaccine efficacy is preserved after up to 5 years of frozen storage, supporting the long-term stability of HK-fbp1 preps.

**FIG 3.**
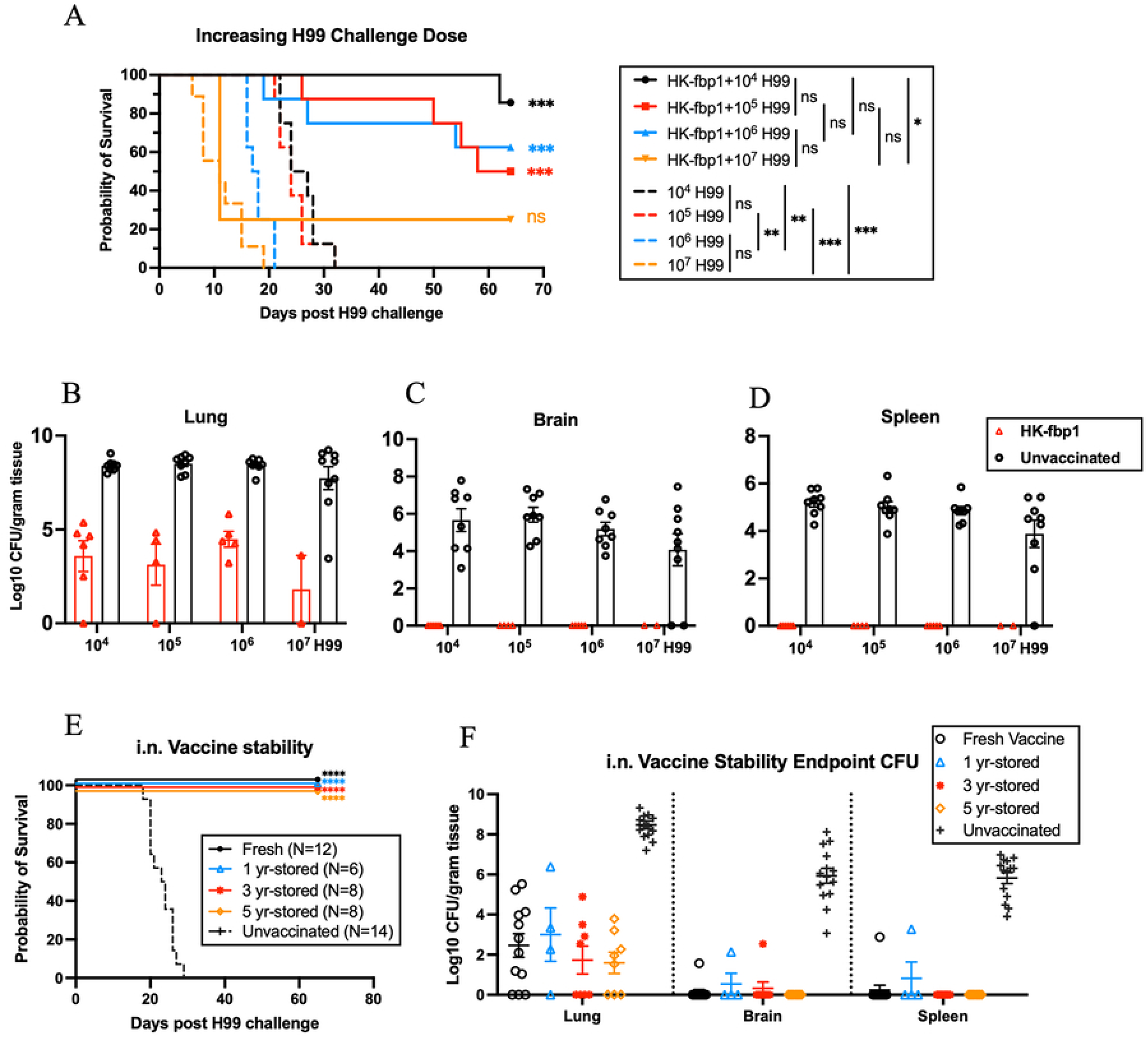
HK-fbp1 vaccination protects mice against a wide range of challenge doses and retains efficacy after long-term frozen storage. (A) Survival curves of mice intranasally vaccinated twice with 5 × 10⁷ HK-fbp1 cells and challenged by infection with 10⁴, 10⁵, 10⁶, or 10⁷ live wild-type H99 cells. Vaccination and challenge were performed as described for Fig. 1B. Unvaccinated mice challenged with the same doses are shown as controls. Seven to 9 BALB/c mice were used for each group. *, *P* < 0.05; ***, *P* < 0.001 (determined by log rank Mantel-Cox test). Median survival times and 95% confidence intervals for all groups are given in Table S1. (B to D) Fungal burden in the lungs (B), brains (C), and spleens (D) of the surviving mice at the end of the experiment (n = 2 to 5 per vaccinated group and 8 to 9 per unvaccinated group). The fungal burden of unvaccinated mice at the humane endpoint is shown as a control. (E) Survival curves of mice immunized twice intranasally with 5 × 10⁷ HK-fbp1 cells prepared fresh or stored at −20°C for 1, 3, or 5 years and challenged by infection with 10⁴ wild-type H99 cells. Vaccination and challenge were performed as described for Fig. 1B. Unvaccinated mice are shown as a control. Six to 14 BALB/c mice were used for each group. *, *P* < 0.05 (determined by log rank Mantel-Cox test). (F) Fungal burden in the lungs, brains, and spleens of the surviving mice at the end of the experiment (n = 4 to 14 per group). The fungal burden of unvaccinated mice at the humane endpoint is shown as a control. Groups were compared by one-way ANOVA with Tukey’s multiple-comparison test within each tissue; comparisons without an asterisk are not significant.

### Supplementation of CpG adjuvant ODN restores robust protection at a suboptimal HK-fbp1 dose

Since protection by HK-fbp1 depends on type 1 immunity (13), we hypothesized that addition of an adjuvant could alleviate the need of using high HK-fbp1 inoculum to achieve full protection in our intranasal model. To test this, we incorporated the Th1-promoting adjuvant CpG ODN2395 into the HK-fbp1 formulation. Using our standard two-dose intranasal regimen (Fig. 1B), the inclusion of ODN allowed the vaccine inoculum to be reduced five-fold, from 5 × 10⁷ to 1 × 10⁷ cells. Whereas 10⁷ HK-fbp1 alone protected only 9% of mice, 10⁷ HK-fbp1 formulated with ODN provided a strong protection with ∼73% survivors, comparable to the 100% protection conferred by 5 × 10⁷ HK-fbp1, while ODN alone did not show any protection and was indistinguishable from unvaccinated controls (Fig. 4A). Mice immunized with 5 × 10⁷ HK-fbp1 or with 10⁷ HK-fbp1 plus ODN lost weight transiently after both immunization on day 0 and 30, whereas weight fluctuation was milder in mice given 10⁷ HK-fbp1 and minimal in mice given ODN alone (Fig. S3A). After H99 challenge, mice immunized with 5 × 10⁷ or 1 × 10⁷ HK-fbp1/ODN gained weight steadily, whereas mice given ODN or left unvaccinated lost weight progressively and succumbed to infection within 5 weeks (Fig. 4A and B). In the 10⁷ HK-fbp1 group, two mice showed prolonged survival but marked weight loss, and one of these died at 6 weeks (Fig. 4A and B).

**FIG 4.**
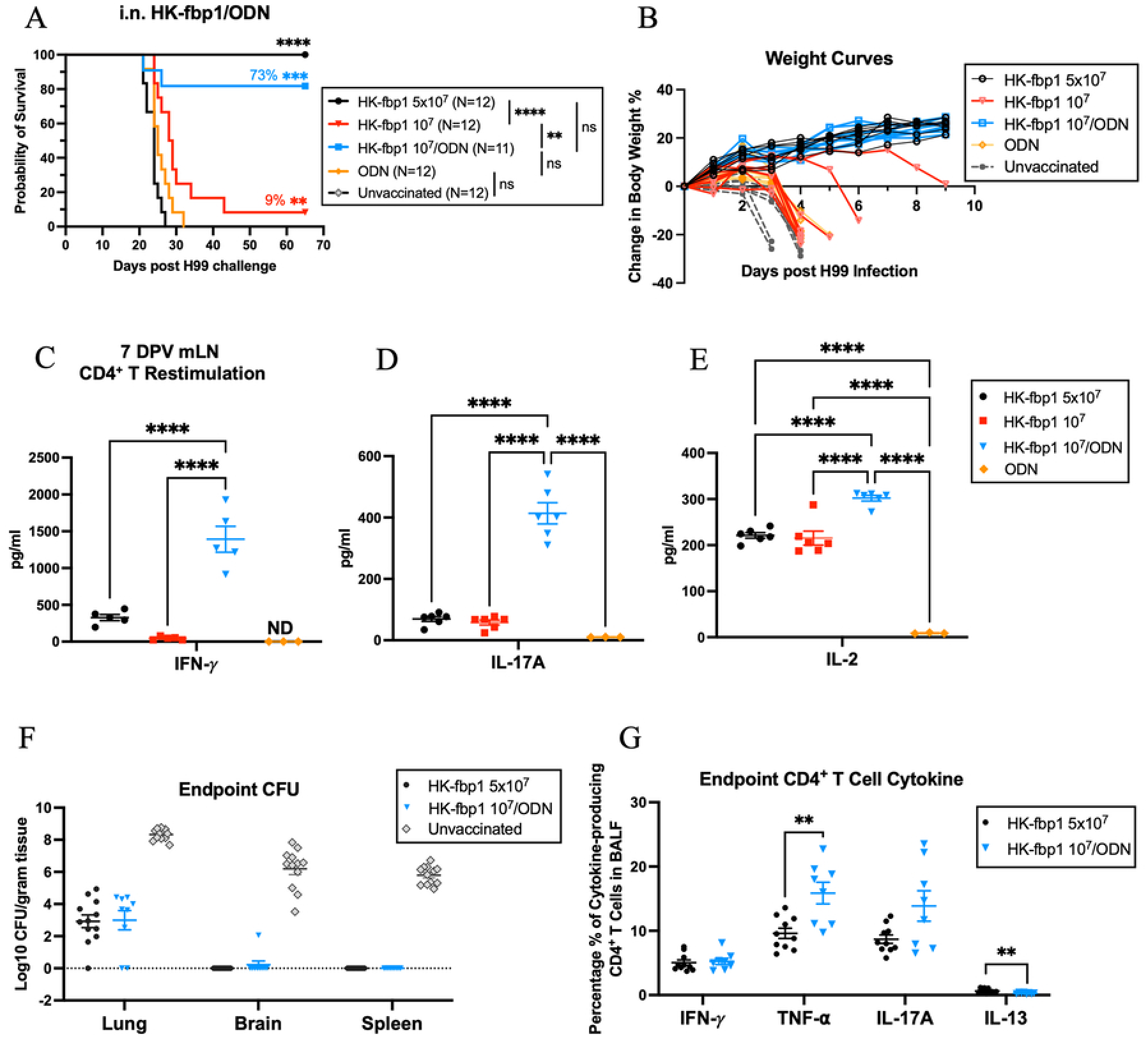
CpG ODN adjuvantation reduces the HK-fbp1 dose required for protection and enhances antigen-specific T cell responses. Mice were vaccinated with 5 × 10⁷ HK-fbp1 cells, 10⁷ HK-fbp1 cells, 10⁷ HK-fbp1 cells formulated with 20 μg CpG ODN 2395 (HK-fbp1/ODN), or CpG ODN 2395 alone. Vaccination and challenge were performed as described for Fig. 1B. (A) Survival curves. Unvaccinated mice are shown as a control. Eleven to 12 BALB/c mice were used for each group. Statistical analysis of survival rates was determined by log rank Mantel-Cox test. **, P < 0.01; ***, P < 0.001; ****, P < 0.0001. (B) Changes in mouse body weight after challenge infection. All living mice for each group were weighed and presented weekly. (C to E) Antigen-specific production of IFN-γ (C), IL-17A (D), and IL-2 (E) by mediastinal lymph node (mLN) CD4⁺ T cells. CD4⁺ T cells were isolated from the mediastinal lymph nodes at 7 days post vaccination (DPV) and cocultured with naive splenic antigen-presenting cells in the presence or absence of H99 antigen. Cytokine levels were compared by one-way ANOVA with Tukey’s multiple-comparison test within each cytokine. ****, P < 0.0001. ND, not detected. (F) Fungal burden in the lungs, brains, and spleens at the end of the experiment (n = 10 to 12 per group). The fungal burden of unvaccinated mice at the humane endpoint is shown as a control. (G) Cytokine production profile of CD4⁺ T cells in BALF at the end of the experiment (n = 8 to 10 per group). Fungal burdens and cytokine levels were compared by one-way ANOVA with Tukey’s multiple-comparison test within each tissue or cytokine. **, P < 0.01; ****, P < 0.0001. Comparisons without an asterisk are not significant.

To characterize the vaccine-induced airway response, we measured antigen-specific cytokine responses at 7 days post vaccination (DPV). CD4⁺ T cells were isolated from the lung-draining mediastinal lymph nodes (mLN) of vaccinated mice and cocultured with naive splenic antigen-presenting cells in the presence of H99 antigen. CD4⁺ T cells from mice immunized with 10⁷ HK-fbp1/ODN produced the highest levels of antigen-specific IFN-γ, IL-17A, and IL-2, followed by those immunized with 5 × 10⁷ HK-fbp1 (Fig. 4C to E). In mice receiving ODN alone, IFN-γ was undetectable and IL-17A and IL-2 remained at baseline. Responses in mice immunized with 10⁷ HK-fbp1 alone were comparable to those in the ODN-alone group, except for significantly higher IL-2 in all HK-fbp1 containing formulations (Fig. 4C to E). The proportion of CD4⁺ T cells in BALF did not differ between groups at 7 DPV (Fig. S3B). Cytokine production by BALF CD4⁺ and CD8⁺ T cells, measured without antigen restimulation, showed a similar Th1/Th17 pattern: cytokine responses were elevated in mice immunized with 5 × 10⁷ HK-fbp1 or 10⁷ HK-fbp1/ODN, with no significant difference between these two groups except for higher IFN-γ from CD8⁺ T cells induced by 10⁷ HK-fbp1/ODN, and remained low in mice receiving 10⁷ HK-fbp1 or ODN alone (Fig. S3C and D). At the end of the experiment, mice immunized with 5 × 10⁷ HK-fbp1 or 10⁷ HK-fbp1/ODN retained low-to-moderate pulmonary fungal burdens relative to unvaccinated controls, most had cleared the brain, and no colonies were recovered from the spleen (Fig. 4F). IFN-γ and IL-17A production by BALF T cells again did not differ between mice immunized with 5 × 10⁷ HK-fbp1 and those immunized with 10⁷ HK-fbp1/ODN, whereas TNF-α production was higher and IL-13 production lower in the 10⁷ HK-fbp1/ODN group (Fig. 4G). Together, these results demonstrate that ODN enhances the protective efficacy of intranasal HK-fbp1 vaccination, allowing a five-fold reduction in vaccine inoculum while maintaining robust protection against *C. neoformans*. This improvement is accompanied by strong antigen-specific Th1/Th17 responses and sustained control of fungal infection.

### HK-fbp1 retains superior protective efficacy over HK-H99 during CpG ODN adjuvanted vaccination in both intranasal and intramuscular models

Having established the protection superiority of HK-fbp1 over HK-H99 in both prophylactic and therapeutic intranasal settings, we next tested whether this difference was maintained when the two heat-killed strains were administered with CpG ODN 2395. Mice were immunized twice intranasally with 10^7^ HK-fbp1 or HK-H99 cells together with 20 μg ODN per dose, and challenged intranasally with 1 x 10^4^ H99 cells. At this dose, HK-fbp1 alone without adjuvant offers minimal/no protection (Fig. 4A) (13), ODN addition enhanced the efficacy of HK-fbp1 (75% survival) but not that of HK-H99 (30%) (Fig. 5A). HK-H99 immunized mice lost ∼14% of their initial body weight by 3 weeks post infection (WPI), whereas HK-fbp1 immunized mice had lost only less than 4% at that time and reached their lowest point a week later before regaining weight (-9% at 4 WPI; Fig. 5B). At the study endpoint, fungal burdens in the lung, brain, and spleen of survival mice were comparable between the two immunized groups but significantly lower than in unvaccinated controls (Fig. 5C). Immune profiling of BALF at the study endpoint showed a comparable pattern of Th1 responses by CD4⁺ and CD8⁺ T cells in HK-H99 and HK-fbp1 immunized mice (Fig. 5D).

**FIG 5.**
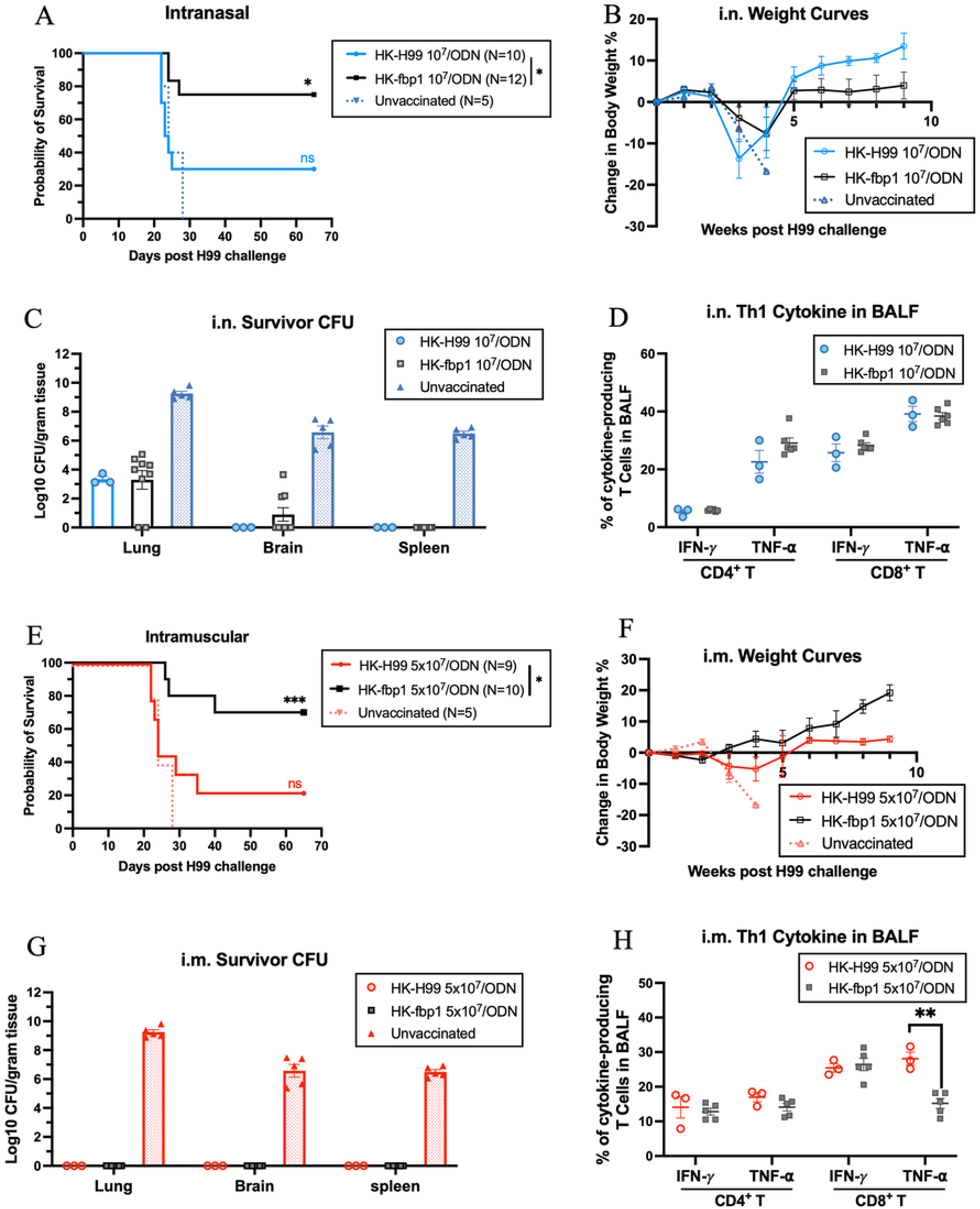
CpG ODN supplementation does not compensate for the weaker protection conferred by HK-H99. (A to D) Mice were immunized twice intranasally, 4 weeks apart, with 10⁷ HK-fbp1 or HK-H99 cells together with 20 μg CpG ODN 2395 per dose and challenged by infection with 10⁴ H99 cells 2 weeks after the boost. Unvaccinated mice are shown as a control. (A) Survival curves. Ten to 12 BALB/c mice were used for each experimental group, and 5 for the control group. Statistical analysis of survival rates was determined by log rank Mantel-Cox test. *, P < 0.05. (B) Changes in mouse body weight after H99 challenge. All living mice for each group were weighed and presented weekly. (C) Fungal burden in the lungs, brains, and spleens of the surviving mice at the end of the experiment. The fungal burden of unvaccinated mice at the humane endpoint is shown as a control. (D) Th1 cytokine production by BALF T cells from the surviving mice shown in panel C at the end of the experiment (n = 3 to 6 per group). Fungal burdens and cytokine levels were compared by Mann-Whitney test within each tissue or cytokine. Comparisons without an asterisk are not significant. (E to H) Mice were immunized twice intramuscularly, 4 weeks apart, with 5 × 10⁷ HK-fbp1 or HK-H99 cells together with 20 μg CpG ODN 2395 per dose and challenged intranasally with 10⁴ H99 cells. (E) Survival curves. Nine to 10 BALB/c mice were used for each experimental group, and 5 for the control group. Statistical analysis of survival rates was determined by log rank Mantel-Cox test. *, P < 0.05; ***, P < 0.001. (F) Changes in mouse body weight after H99 challenge. (G) Fungal burden in the lungs, brains, and spleens of the surviving mice at the end of the experiment. The fungal burden of unvaccinated mice at the humane endpoint is shown as a control. (H) Th1 cytokine production by BALF T cells from the surviving mice shown in panel G at the end of the experiment (n = 3 to 5 per group). Fungal burdens and cytokine levels were compared by Mann-Whitney test within each tissue or cytokine. Comparisons without an asterisk are not significant. **, P < 0.01. The same unvaccinated control group is shown for both the intranasal and intramuscular experiments in this figure, as the two experiments were performed concurrently.

In recent work, we characterized the protection conferred by intramuscular HK-fbp1/ODN vaccination and its immunological requirements, however, HK-H99 was not evaluated in the presence of ODN (Y. Zhang et al., submitted for publication). To determine whether the advantage of HK-fbp1 is retained when the vaccine is delivered by a parenteral route, mice were immunized twice intramuscularly with 5 × 10⁷ HK-fbp1/ODN or HK-H99/ODN and challenged intranasally with H99. Similar to the intranasal setting, the adjuvant enhanced the protection of HK-fbp1 but did not compensate for the weaker protection conferred by HK-H99: HK-fbp1/ODN protected 70% of mice compared with 30% for HK-H99/ODN (Fig. 5E). Consistent with the stronger protection, HK-fbp1/ODN immunized mice gained weight more steadily (Fig. 5F). Notably, no fungal cells were recovered from any organs examined at the study endpoint after intramuscular immunization with either preparation, suggesting this combination regimen could potentially provide sterilizing immunization (Fig. 5G). T cell profiling did not distinguish the two formulations, with comparable BALF Th1 cytokine production by both subsets (Fig. 5H). Together, these results show that in both intranasal and intramuscular settings with supplementation of ODN, HK-fbp1 is also outperforms HK-H99. Within each route, mice that survived either immunization controlled the infection to a similar extent, and no difference in Th1 cytokine production by two vaccine formulations under both immunization routes was observed, indicating that the two formulations differ primarily in the likelihood of establishing protective immunity rather than in the quality of the control achieved once it is established. Our data also showed that intramuscular immunization completely cleared fungal infection in survival mice, suggesting this immunization strategy can potentially reach sterilizing immunization. On the other hand, intranasally vaccinated mice that survived H99 challenge retained residual pulmonary burdens.

### Intramuscular vaccination of HK-fbp1/ODN confers complete and durable protection

We next defined the protection thresholds of this adjuvanted vaccination strategy, including the minimum antigen dose, the minimum number of doses, and the duration of protection. We first titrated the minimal HK-fbp1 cell numbers required to confer protection in the presence of ODN adjuvant by administering 10⁷, 10⁶, or 10⁵ cells together with 20 μg ODN under the two-dose regimen (Fig. 6A). Complete protection was retained when 10⁷ cells plus ODN was used, whereas neither 10⁶ nor 10⁵ cells conferred any protection (Fig. 6B). Protection thus declined abruptly between 10⁷ and 10⁶ cells, indicating a narrow antigen-dose threshold for this route (Fig. 6B). We then asked whether a single immunization would be sufficient to provide protection. Mice received one intramuscular dose of 5 × 10⁷ or 10⁷ HK-fbp1 cells plus ODN and were challenged with 10^4^ H99 at day 30 post vaccination (Fig. 6C). We found that a single 5 × 10⁷ HK-fbp1/ODN dose conferred complete protection, whereas a single 10⁷ dose protected 37.5% of mice, still a significant improvement than unvaccinated controls (Fig. 6D). Protection after a single dose immunization was thus also dose dependent, mirroring the dose dependence observed with the intranasal route. Finally, we tested the durability of single-dose protection (Fig. 6E). Mice challenged 3 months after a single 5 × 10⁷ HK-fbp1/ODN intramuscular immunization remained fully protected, as did those challenged at one month post vaccination (Fig. 6F). Together, these results define the protection range of the intramuscular vaccine formulation: under a two-dose regimen, protection required at least 10⁷ cells plus ODN and was abruptly lost below this number, whereas a single 5 × 10⁷ HK-fbp1/ODN dose was sufficient to confer complete protection that persisted for at least 3 months, indicating that our vaccine mediated immune protection is long lasting.

**FIG 6.**
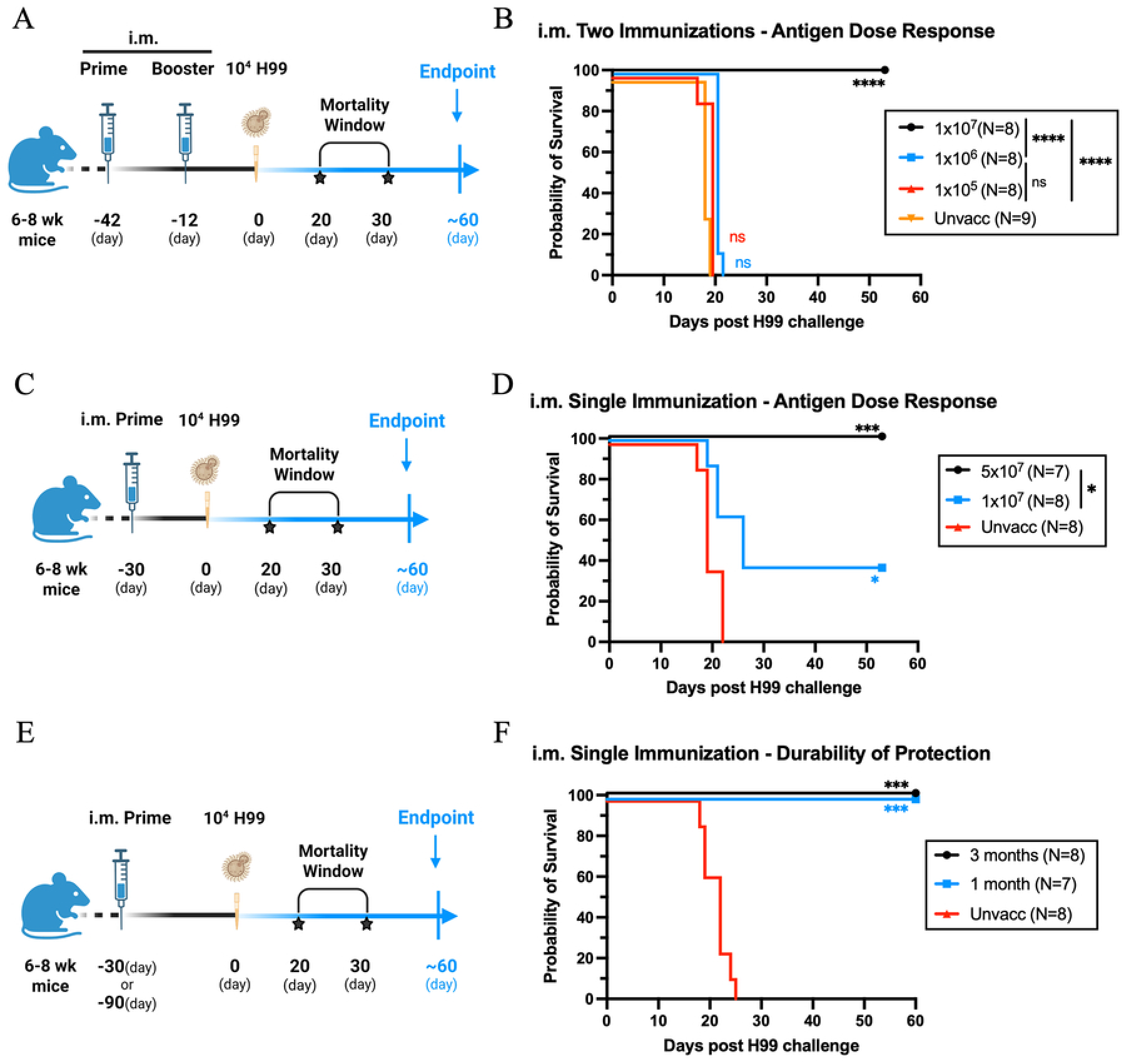
Intramuscular immunization(s) of adjuvanted HK-fbp1 confers strong and durable protection. (A) Immunization and challenge schedule. C57BL/6 mice were immunized intramuscularly with 10⁷, 10⁶, or 10⁵ HK-fbp1 cells together with 20 μg CpG ODN 2395 per dose and challenged intranasally with 10⁴ H99 cells. Unvaccinated mice are shown as a control. (B) Survival curves of mice immunized and challenged as described in panel A. Eight to 9 C57BL/6 mice were used for each group. (C) Immunization and challenge schedule. C57BL/6 mice immunized once with 5 × 10⁷ or 1× 10⁷ HK-fbp1 cells and challenged 30 DPV. (D) Survival curves of mice immunized and challenged as described in panel C. Eight to 9 mice were used for each group. ****, P < 0.0001. (E) Immunization and challenge schedule. C57BL/6 mice were immunized once with 5 × 10⁷ HK-fbp1 cells and challenged 30 days or 90 days later. Seven to 8 C57BL/6 mice were used for each group. *, P < 0.05; ***, P < 0.001. (F) Survival curves of mice immunized and challenged as described in panel A. Seven to 8 C57BL/6 mice were used for each group. The schematic was created with BioRender (A, C, E). Statistical analysis of survival rates was determined by log rank Mantel-Cox test (B, D, F). *, P < 0.05; ***, P < 0.001; ****, P < 0.0001.

## Discussion

In previous studies we uncovered that infection with *fbp1*Δ mutant cells could induce a superior Th1/Th17 immune response as compared to H99 infection. Moreover, we previously determined that *fbp1*Δ mutant heat-killed cells remain highly immunogenic and can protect mice from challenge with highly virulent wild type *C. neoformans* strain H99 and *C. gattii* strain R265 (12, 13). Intriguingly, when used in a high dose, immunization with heat-killed H99 has also been reported to provide certain level of fungal control and protection against *C. neoformans* infection (16, 17). The HK-H99 whole cell mediated protective immunity is dependent on the culture conditions that induced different cell surface composition (18). Although we used HK-H99 as a control in our previous studies (12), a systematic comparison of the protection and immune activation conferred by HK-fbp1 and HK-H99 had not been performed before. The findings of the current study indicate that there is a clear protective advantage conferred by HK-fbp1 over HK-H99. Our results suggest that HK-fbp1vaccine is more potent than HK-H99 in a per cell basis although this protective ability is not exclusive to HK-fbp1. Although HK-fbp1 provided stronger protection under most conditions tested, HK-H99 was not inert: at the highest antigen dose, it conferred measurable protection. Among animals that survived, fungal burdens were comparable between the two preparations, although such comparisons may be affected by selection bias, particularly when survival differs markedly between vaccination groups. Our findings also suggest that HK-fbp1 is more effective than HK-H99 at inducing protective immunity. Because Fbp1 regulates multiple fungal cellular processes, its deletion may alter antigen abundance, cell surface composition, or accessibility of immunostimulatory components before heat inactivation (12). We hypothesize that the *fbp1*Δ cells overproduce various antigenic factors as compared to H99 thus becoming able to induce more robust protective immunity. In previous studies we identified several factors that contribute to *fbp1*Δ mediated immune stimulation, including capsule components and the mannoprotein Cig1 (20). Further characterization of the antigenic and structural differences between these preparations will therefore be important for understanding the basis of enhanced HK-fbp1 vaccine potency and may offer an opportunity to further optimize the vaccine regimen by including these individual immunogenic factors in the vaccine formulation. Similar antigen identification efforts have been reported in other studies with the goal to develop a subunit based antifungal vaccine (21, 22). Differences in T-cell response and differentiation may provide additional insight into this discrepancy. Compared with HK-H99, HK-fbp1 vaccination was associated with greater representation of CD4⁺ T cells in the lung compartment, together with higher proportions of Th1 cytokine production and increased expression of activation markers. Future studies comparing antigen-specific T-cell phenotypes and effector functions induced by these two vaccines may help clarify the immunological basis of their different protective efficacy.

Our findings also highlight the importance of distinguishing vaccine-induced protection from the capacity to withstand overwhelming fungal exposure. Although HK-fbp1 remained protective against substantially increased challenge inoculum, survival declined sharply at 10^7^ CFU inoculum tested. Interestingly, the vaccinated mice that succumbed at this dose showed survival kinetics similar to those of unvaccinated controls. One possible explanation is that successful protection requires early containment of fungal proliferation before fungal burden exceeds the capacity of vaccine-primed immune responses to control the spread of infection. The polysaccharide capsule of *C. neoformans*, which contributes to immune evasion and fungal persistence, may further complicate clearance under high-burden conditions (23). However, under the experimental conditions used, we are unable to determine whether protection failed during initial fungal containment or during subsequent control of established infection. Given the importance of innate immunity in early antifungal defense (15), longitudinal analysis of fungal burden and innate immune responses following high-dose challenge would help define when and how protective immunity is established or fails. Importantly, these high-inoculum experiments establish the robustness and thresholds of vaccine-induced protection rather than directly reproducing natural infectious exposure. It is also exciting to observe that long-term (5 years) frozen storage did not compromise protective efficacy of HK-Fbp1 vaccine preparations under the conditions tested, suggesting the vaccine formulation is highly stable.

Adjuvants are widely used to enhance vaccine immunogenicity, protective efficacy and durability (24-26). Several adjuvants have been reported in fungal vaccine development, such as cationic adjuvant formulation 01 (CAF01) and alum (27, 28). We are excited to find that adding CpG adjuvant ODN 2395 significantly improved our HK-fbp1 vaccine. Addition of ODN 2395 enabled enhanced protection, and allowed us to identify additional immunization routes, including the most commonly used intramuscular injection method. From a translational perspective, the benefit of CpG ODN adjuvantation extends beyond increasing the magnitude of protection. The vaccine antigen dose threshold required for protection after intranasal delivery was significantly reduced by adjuvantation with ODN 2395. Notably, all HK-fbp1-containing formulations elicited antigen-specific IL-2 recall responses, whereas ODN alone did not, suggesting that HK-fbp1 antigen can prime antigen-specific CD4⁺ T cell memory responses without CpG adjuvantation. However, the low-dose (10⁷ cells) HK-fbp1 formulation induced minimal IFN-γ and IL-17A responses and provided little protection despite its detectable IL-2 response. In contrast, the addition of CpG ODN markedly enhanced IFN-γ and IL-17A production, accompanied by substantially improved survival. These findings suggest that antigen-specific T-cell responsiveness alone is insufficient for protection and that CpG adjuvantation promotes Th1/Th17-associated effector responses that correlate with protective immunity. Additionally, CpG ODN enables effective intramuscular delivery of HK-fbp1. Intranasal vaccinated mice exhibited transient but sharp weight loss and observable signs of discomfort, whereas intramuscularly vaccinated mice exhibited steady weight gain (data not shown), suggesting an additional practical advantage of intramuscular administration. The strong dose dependence observed in both vaccination models also emphasizes the need for standardized vaccine preparations. The successful single-dose regimen in the intramuscular setting suggests robust potency and that repeated immunization may not be necessary to achieve strong protection in immunocompetent hosts. However, the efficacy of the optimized intramuscular formulation in hosts with advanced cellular immunodeficiency remains an important unresolved question.

Overall, in this study, we have defined several characteristics of our HK-fbp1 vaccine that are important for its further development. We found our vaccine candidate is stable with superior protectivity than the heat-killed wild type cells. Its CpG-adjuvanted version (HK-fbp1/ODN) works in a single dose in an intramuscular immunization model with long lasting immune protection. This study supports a model in which the protective efficacy of inactivated cryptococcal vaccines depends not only on the fungal strain used for vaccine preparation but also on antigen dose, adjuvant formulation and administration route. These findings provide a rationale for investigating the antigenic properties of HK-fbp1 and the functional contributions of vaccine-induced T-cell populations while advancing a more practical formulation for future antifungal vaccine development.

## Materials and Methods

### Ethics statement on animal use

Female mice weighing 20–25 g were used in all experiments. Wild-type (WT) BALB/c and C57BL/6 mice were purchased from The Jackson Laboratory. All animal experiments were carried out at the Rutgers University Newark-Public Health Research Institute (PHRI) animal facility in accordance with biosafety level 2 (BSL-2) procedures and protocols approved by the Rutgers University Institutional Animal Care and Use Committee (IACUC) and Institutional Biosafety Committee (protocol no. 999901066). All animal care and experimental procedures complied with the Animal Welfare Act and the Public Health Service (PHS) Policy on Humane Care and Use of Laboratory Animals.

### Vaccination strategies

*C. neoformans fbp1*Δ mutant strain or wild type H99 was heat-killed following a previously described procedure (13). In brief, *C. neoformans* cells were recovered on YPD agar with corresponding selective marker, inoculated into liquid YPD medium, and grown overnight at 30°C with shaking at 225 rpm. Cells were collected by centrifugation at 600-800 × g for 5 min at room temperature (RT), washed 2-3 times in 1× PBS, and counted with a hemocytometer. Washed cells were heat killed at 75-77°C for 90 min and resuspended in 1× PBS for immunization. Loss of viability and concentration confirmation was confirmed by plating an aliquot of live and heat-killed suspension on YPD agar. CpG ODN 2395 (InvivoGen, CAT# vac-2395-1) was reconstituted according to the manufacturer’s instructions. For vaccine preparation, heat-killed *C. neoformans* cells were suspended in 50 μl of 1× PBS. To formulate ODN-adjuvanted HK-fbp1 vaccines (HK-fbp1/ODN), 10 μl of supernatant was removed from the cell suspension and replaced with an equal volume of ODN 2395 stock solution (2 mg/ml). For the ODN-only control, 10 μl ODN 2395 stock solution (2 mg/ml) was diluted with 40 μl of sterile, endotoxin-free physiological water. All vaccine preparations were thoroughly mixed immediately before administration. Vaccination strategies and timelines are specified in the corresponding figure legends.

### Animal infections with *C. neoformans*

To prepare fungal cells for infection, *C. neoformans* H99 cells grown overnight in YPD broth were washed two to three times with 1× PBS, and yeast cell concentrations were determined using a hemocytometer. Mice were anesthetized with a mixture of ketamine (12.5 mg/ml) and xylazine (1 mg/ml) before intranasal infection with 1 × 10⁴, 1 × 10⁵, 1 × 10⁶, or 1 × 10⁷ live H99 cells in a total volume of 50 μl. Following infection, mice were weighed daily and monitored twice daily for signs of disease progression, including weight loss, gait abnormalities, labored breathing, and ruffled fur. Animals exhibiting severe distress or becoming moribund were euthanized by CO₂ inhalation. Survival data were plotted over time, and differences between groups were analyzed using the log-rank (Mantel–Cox) test in GraphPad Prism version 10.5 (GraphPad Software). P values of less than 0.05 were considered statistically significant.

### Fungal burdens determination

To determine fungal burdens, lungs, brains, and spleens were collected from infected mice at the indicated time points and individually homogenized in 3 ml of cold 1× PBS for 1 min using a Fisher brand Homogenizer 150 (Fisher Scientific, MA). Tissue homogenates were serially diluted and plated onto YPD agar supplemented with ampicillin and chloramphenicol. Colony-forming units (CFU) were counted after incubation at 30°C for 2 to 3 days.

### Lung tissue processing and flow cytometry

Single cell suspensions of pulmonary cells were prepared for flow cytometric analysis as previously described (29, 30). In brief, lung tissues were minced in 5 ml of 1× PBS containing 3 mg/ml collagenase type IV (Worthington) and incubated at 37°C for 45 min, followed by washing with 1× PBS. Following digestion, residual red blood cells (RBCs) were removed using RBC lysis buffer (155 mM NH₄Cl and 10 mM NaHCO₃, pH 7.2). The resulting lung single-cell suspensions were used for flow cytometric analysis. Cells were stained with the following antibodies to identify different immune cell populations: monocytes, CD45 (30-F11, BUV395), CD11b (M1/70, PerCP-Cy5.5), and Ly6C (AL-21, PE); monocyte-derived dendritic cells (Mo-DCs), CD45 (30-F11, BUV395), CD11b (M1/70, PerCP-Cy5.5), Ly6C (AL-21, PE), CD11c (HL3, BV510), and MHC class II I-A/I-E (M5/114.15.2, BV711); neutrophils, CD45 (30-F11, APC-Cy7), CD11b (M1/70, PerCP-Cy5.5), Ly6C (AL-21, PE), and Ly6G (1A8, APC); CD4⁺ T cells, CD45 (30-F11, BUV395) and CD4 (RM4-5, BV421, and/or RM4-4, FITC); and CD8⁺ T cells, CD45 (30-F11, BUV395), CD8α (53-6.7, BV711), CD62L (MEL-14, APC), and CD44 (IM7, PE). All antibodies were purchased from BD Biosciences. Samples were acquired using a BD LSRFortessa flow cytometer and analyzed with FlowJo software (Tree Star, Inc.).

### Intracellular cytokine staining of T cells harvested in BALF and Flow Cytometry

To assess host immune responses, bronchoalveolar lavage fluid (BALF) was collected at the indicated time points after infection. Following euthanasia, a catheter was inserted into the trachea, and the airways were lavaged five times with approximately 1 ml of 1× PBS per wash (5 ml total). After red blood cell (RBC) lysis, BALF cells were transferred to 96-well round-bottom plates and restimulated for 5 to 6 h with BD Leukocyte Activation Cocktail containing BD GolgiPlug (BD Biosciences), according to the manufacturer’s instructions. Cells were then stained with fluorescently labeled antibodies against surface markers and fixed overnight in 1% paraformaldehyde. Following fixation, cells were permeabilized with 1× BD Perm/Wash buffer (BD Biosciences) and stained for intracellular cytokines for 30 min on ice using antibodies diluted in the same buffer. Samples were washed and analyzed by flow cytometry. Antibodies used for surface staining included Thy1.2 (53-2.1, PE-Cy7), CD4 (RM4-5, BV421), and CD8α (53-6.7, PerCP-Cy5.5). Intracellular cytokine staining was performed using antibodies against IFN-γ (XMG1.2, PE), IL-17A (eBio17B7, APC), TNF-α (MP6-XT22, BV711), and IL-13 (eBio13A, FITC). All antibodies and reagents were obtained from BD Biosciences, except the IL-17A and IL-13 antibodies, which were purchased from eBioscience.

### CD4^+^ T cell isolation and CD4^+^ T cell recall response

Antigen-presenting cells (APCs) were prepared from the spleens of syngeneic, uninfected donor mice. Splenic tissues were gently disrupted using the frosted ends of two glass slides to release leukocytes, followed by T-cell depletion through antibody- and complement-mediated lysis. Briefly, splenic cells were incubated with anti-Thy1.2 antibodies (BioXCell, CAT# BE00066) and rabbit complement (Low-Tox; Cedarlane Labs, CAT# CL3331) at 37°C for 1 h. Cell suspensions were then passed through a 70-μm nylon mesh. Lung-draining mediastinal lymph nodes (mLNs) were collected in 10 ml of 1× PBS and mechanically dissociated using the frosted ends of two glass slides to obtain single-cell suspensions. Samples from individual mice within each group were pooled for CD4⁺ T-cell isolation. CD4⁺ T cells were purified by negative selection using a CD4⁺ T-cell isolation kit (Miltenyi Biotec, Inc.; CAT# 130-104-454) according to the manufacturer’s instructions. The purity of isolated CD4⁺ T cells was consistently greater than 90%, as determined by flow cytometry. Purified CD4⁺ T cells (2 × 10⁵) were cocultured with T-cell-depleted splenic APCs (3 × 10⁵) in RPMI medium (Corning, CAT# 15-040-CV) supplemented with 10% fetal bovine serum (FBS; Cytiva, CAT# SH30910.03), penicillin-streptomycin (2,200 U/ml; Gibco, CAT# 15140-122), and gentamicin sulfate (1 mg/ml). To assess *Cryptococcus*-specific CD4⁺ T-cell recall responses, cocultures were stimulated with bead-beaten H99 yeast cells prepared using an MP FastPrep Grinder as a source of fungal antigens, at an APC-to-yeast ratio of 1:1.5. Voriconazole was added at a final concentration of 0.5 mg/ml to prevent fungal outgrowth during incubation. After 72 h at 37°C with 5% CO₂, culture supernatants were collected, and IL-2, IL-17A, and IFN-γ levels were measured using mouse uncoated ELISA kits (Invitrogen) according to the manufacturer’s instructions.

## Data Availability

All source data supporting the findings of this study are provided in the Supplementary Data.

## Statistics and Reproducibility

GraphPad Prism was used for statistical analyses of survival, fungal burden, flow cytometry, and other experimental datasets. Data are presented as mean ± SD unless otherwise indicated. Each data point represents an independent biological sample or experimental replicate, unless otherwise stated. Technical replicates, when performed, were averaged and were not treated as independent observations. Exact sample sizes (N) and statistical tests are provided in the corresponding figure or figure legends. All statistical tests were two-sided, and P < 0.05 was considered statistically significant. No data were excluded unless otherwise stated. Experiments were independently repeated at least two to three times unless otherwise specified.

## Acknowledgments

This study is supported by NIH grants AI169769 and AI141368 to CX and AR. The Xue lab is also supported by NIH grants AI123315 and AI155647. AR holds an Investigators in the Pathogenesis of Infectious Disease Award from the Burroughs Welcome Fund.

## Author contributions

Y.Z, A.R. and C.X. developed hypothesis and study design. Y.Z. and C.X. wrote the first and successive drafts of the manuscript. Y.Z, Y.W, K.W, L.G, S.P, A.L contributed to interpretation of data. Y.Z, Y.W, K.W, L.G performed experiments. A.R. and C.X. obtained funding.

## Reference

1. Brown GD, Denning DW, Levitz SM. Tackling human fungal infections. Science. 2012;336(6082):647. doi: 10.1126/science.1222236. PubMed PMID: 22582229.

2. Brown GD, Denning DW, Gow NA, Levitz SM, Netea MG, White TC. Hidden killers: human fungal infections. Sci Transl Med. 2012;4(165):165rv13. doi: 10.1126/scitranslmed.3004404. PubMed PMID: 23253612.

3. Rajasingham R, Govender NP, Jordan A, Loyse A, Shroufi A, Denning DW, et al. The global burden of HIV-associated cryptococcal infection in adults in 2020: a modelling analysis. Lancet Infect Dis. 2022;22(12):1748–55. Epub 20220829. doi: 10.1016/S1473-3099(22)00499-6. PubMed PMID: 36049486; PubMed Central PMCID: PMC9701154.

4. Oliveira LVN, Wang R, Specht CA, Levitz SM. Vaccines for human fungal diseases: close but still a long way to go. NPJ Vaccines. 2021;6(1):33. Epub 20210303. doi: 10.1038/s41541-021-00294-8. PubMed PMID: 33658522; PubMed Central PMCID: PMC7930017.

5. Rivera A, Lodge J, Xue C. Harnessing the Immune Response to Fungal Pathogens for Vaccine Development. Annu Rev Microbiol. 2022;76:703–26. Epub 20220627. doi: 10.1146/annurev-micro-041020-111511. PubMed PMID: 35759871; PubMed Central PMCID: PMC11926770.

6. Cutler JE, Deepe GS, Jr., Klein BS. Advances in combating fungal diseases: vaccines on the threshold. Nat Rev Microbiol. 2007;5(1):13–28. Epub 20061211. doi: 10.1038/nrmicro1537. PubMed PMID: 17160002; PubMed Central PMCID: PMC2214303.

7. Van Dyke MCC, Chaturvedi AK, Hardison SE, Leopold Wager CM, Castro-Lopez N, Hole CR, et al. Induction of Broad-Spectrum Protective Immunity against Disparate Cryptococcus Serotypes. Front Immunol. 2017;8:1359. Epub 20171030. doi: 10.3389/fimmu.2017.01359. PubMed PMID: 29163469; PubMed Central PMCID: PMC5670106.

8. Rella A, Mor V, Farnoud AM, Singh A, Shamseddine AA, Ivanova E, et al. Role of Sterylglucosidase 1 (Sgl1) on the pathogenicity of Cryptococcus neoformans: potential applications for vaccine development. Front Microbiol. 2015;6:836. Epub 20150811. doi: 10.3389/fmicb.2015.00836. PubMed PMID: 26322039; PubMed Central PMCID: PMC4531891.

9. Normile TG, Del Poeta M. Three Models of Vaccination Strategies Against Cryptococcosis in Immunocompromised Hosts Using Heat-Killed Cryptococcus neoformans Deltasgl1. Front Immunol. 2022;13:868523. Epub 20220509. doi: 10.3389/fimmu.2022.868523. PubMed PMID: 35615354; PubMed Central PMCID: PMC9124966.

10. Zhai B, Wozniak KL, Masso-Silva J, Upadhyay S, Hole C, Rivera A, et al. Development of protective inflammation and cell-mediated immunity against Cryptococcus neoformans after exposure to hyphal mutants. mBio. 2015;6(5):e01433–15. Epub 20151006. doi: 10.1128/mBio.01433-15. PubMed PMID: 26443458; PubMed Central PMCID: PMC4611043.

11. Upadhya R, Lam WC, Maybruck B, Specht CA, Levitz SM, Lodge JK. Induction of Protective Immunity to Cryptococcal Infection in Mice by a Heat-Killed, Chitosan-Deficient Strain of Cryptococcus neoformans. mBio. 2016;7(3). Epub 20160510. doi: 10.1128/mBio.00547-16. PubMed PMID: 27165801; PubMed Central PMCID: PMC4959652.

12. Masso-Silva J, Espinosa V, Liu TB, Wang Y, Xue C, Rivera A. The F-Box Protein Fbp1 Shapes the Immunogenic Potential of Cryptococcus neoformans. mBio. 2018;9(1). Epub 20180109. doi: 10.1128/mBio.01828-17. PubMed PMID: 29317510; PubMed Central PMCID: PMC5760740.

13. Wang Y, Wang K, Masso-Silva JA, Rivera A, Xue C. A Heat-Killed Cryptococcus Mutant Strain Induces Host Protection against Multiple Invasive Mycoses in a Murine Vaccine Model. mBio. 2019;10(6). Epub 20191126. doi: 10.1128/mBio.02145-19. PubMed PMID: 31772051; PubMed Central PMCID: PMC6879717.

14. Wang Y, Wang K, Rivera A, Xue C. Development of a Heat-Killed fbp1 Mutant Strain as a Therapeutic Agent To Treat Invasive Cryptococcus Infection. Microbiol Spectr. 2023;11(2):e0495522. Epub 20230131. doi: 10.1128/spectrum.04955-22. PubMed PMID: 36719231; PubMed Central PMCID: PMC10101017.

15. Wang K, Espinosa V, Wang Y, Lemenze A, Kumamoto Y, Xue C, et al. Innate cells and STAT1-dependent signals orchestrate vaccine-induced protection against invasive Cryptococcus infection. mBio. 2024;15(10):e0194424. Epub 20240926. doi: 10.1128/mbio.01944-24. PubMed PMID: 39324785; PubMed Central PMCID: PMC11481872.

16. Lin J, Pham T, Hipsher K, Glueck N, Fan Y, Lin X. Immunoprotection against Cryptococcosis Offered by Znf2 Depends on Capsule and the Hyphal Morphology. mBio. 2022;13(1):e0278521. Epub 20220111. doi: 10.1128/mbio.02785-21. PubMed PMID: 35012334; PubMed Central PMCID: PMC8749420.

17. Pham T, Li Y, Watford W, Lin X. Vaccination with a ZNF2(oe) Strain of Cryptococcus Provides Long-Lasting Protection against Cryptococcosis and Is Effective in Immunocompromised Hosts. Infect Immun. 2023;91(7):e0019823. Epub 20230620. doi: 10.1128/iai.00198-23. PubMed PMID: 37338404; PubMed Central PMCID: PMC10353382.

18. Upadhya R, Lam WC, Hole CR, Vasselli JG, Lodge JK. Cell wall composition in Cryptococcus neoformans is media dependent and alters host response, inducing protective immunity. Front Fungal Biol. 2023;4. Epub 20230512. doi: 10.3389/ffunb.2023.1183291. PubMed PMID: 37538303; PubMed Central PMCID: PMC10399910.

19. Kayraklioglu N, Horuluoglu B, Klinman DM. CpG Oligonucleotides as Vaccine Adjuvants. Methods Mol Biol. 2021;2197:51–85. doi: 10.1007/978-1-0716-0872-2_4. PubMed PMID: 32827132.

20. Avina SL, Pawar S, Kadam RN, Rivera A, Xue C. Mannoprotein Cig1 contributes to the immunogenicity of a heat-killed F-box protein Fbp1 Cryptococcus neoformans vaccine model. Infect Immun. 2026;94(1):e0035525. Epub 20251128. doi: 10.1128/iai.00355-25. PubMed PMID: 41313167; PubMed Central PMCID: PMC12797982.

21. Specht CA, Oliveira LVN, Carlson D, Hester MM, Sahu S, Pedersen GK, et al. A protective hybrid protein vaccine informed by mapping major histocompatibility class II epitopes in Cryptococcus neoformans chitin deacetylases that stimulate CD4(+) T-cell responses. mBio. 2026:e0217626. Epub 20261014. doi: 10.1128/mbio.02176-26. PubMed PMID: 42872872.

22. Specht CA, Wang R, Oliveira LVN, Hester MM, Gomez C, Mou Z, et al. Immunological correlates of protection mediated by a whole organism, Cryptococcus neoformans, vaccine deficient in chitosan. mBio. 2024;15(8):e0174624. Epub 20240709. doi: 10.1128/mbio.01746-24. PubMed PMID: 38980038; PubMed Central PMCID: PMC11323574.

23. O’Meara TR, Alspaugh JA. The Cryptococcus neoformans capsule: a sword and a shield. Clin Microbiol Rev. 2012;25(3):387–408. doi: 10.1128/CMR.00001-12. PubMed PMID: 22763631; PubMed Central PMCID: PMC3416491.

24. Reed SG, Orr MT, Fox CB. Key roles of adjuvants in modern vaccines. Nat Med. 2013;19(12):1597–608. Epub 20131205. doi: 10.1038/nm.3409. PubMed PMID: 24309663.

25. Pulendran B, P SA, O’Hagan DT. Emerging concepts in the science of vaccine adjuvants. Nat Rev Drug Discov. 2021;20(6):454–75. Epub 20210406. doi: 10.1038/s41573-021-00163-y. PubMed PMID: 33824489; PubMed Central PMCID: PMC8023785.

26. Coffman RL, Sher A, Seder RA. Vaccine adjuvants: putting innate immunity to work. Immunity. 2010;33(4):492–503. doi: 10.1016/j.immuni.2010.10.002. PubMed PMID: 21029960; PubMed Central PMCID: PMC3420356.

27. Wang R, Oliveira LVN, Hester MM, Carlson D, Christensen D, Specht CA, et al. Protection against experimental cryptococcosis elicited by Cationic Adjuvant Formulation 01-adjuvanted subunit vaccines. PLoS Pathog. 2024;20(7):e1012220. Epub 20240708. doi: 10.1371/journal.ppat.1012220. PubMed PMID: 38976694; PubMed Central PMCID: PMC11257399.

28. Stempinski PR, Ramos Irizarry P, McConnell SA, Liporagi Lopes LC, Rodrigues Dos Santos Junior S, Wear MP, et al. A Cryptococcus neoformans polysaccharide conjugate vaccine made with filtered polysaccharide elicits protective immunity in mice. Fungal Biol. 2025;129(2):101544. Epub 20250125. doi: 10.1016/j.funbio.2025.101544. PubMed PMID: 40023532; PubMed Central PMCID: PMC13316834.

29. Rivera A, Van Epps HL, Hohl TM, Rizzuto G, Pamer EG. Distinct CD4+-T-cell responses to live and heat-inactivated Aspergillus fumigatus conidia. Infect Immun. 2005;73(11):7170–9. doi: 10.1128/IAI.73.11.7170-7179.2005. PubMed PMID: 16239511; PubMed Central PMCID: PMC1273880.

30. Rivera A, Ro G, Van Epps HL, Simpson T, Leiner I, Sant’Angelo DB, et al. Innate immune activation and CD4+ T cell priming during respiratory fungal infection. Immunity. 2006;25(4):665–75. Epub 20061005. doi: 10.1016/j.immuni.2006.08.016. PubMed PMID: 17027299.

